# A multiscale model of the action of a capsid assembly modulator for the treatment of chronic hepatitis B

**DOI:** 10.1101/2024.07.16.603658

**Authors:** Sarafa A. Iyaniwura, Tyler Cassidy, Ruy M. Ribeiro, Alan S. Perelson

**Affiliations:** Theoretical Biology and Biophysics, Theoretical Division, Los Alamos National Laboratory, Los Alamos, NM 87545, USA; School of Mathematics, University of Leeds, Leeds, LS2 9JT, United Kingdom

## Abstract

Chronic hepatitis B virus (HBV) infection is strongly associated with increased risk of liver cancer and cirrhosis. While existing treatments effectively inhibit the HBV life cycle, viral rebound occurs rapidly following treatment interruption. Consequently, functional cure rates of chronic HBV infection remain low and there is increased interest in a novel treatment modality, capsid assembly modulators (CAMs). Here, we develop a multiscale mathematical model of CAM treatment in chronic HBV infection. By fitting the model to participant data from a phase I trial of the first-generation CAM vebicorvir, we estimate the drug’s dose-dependent effectiveness and identify the physiological mechanisms that drive the observed biphasic decline in HBV DNA and RNA, and mechanistic differences between HBeAg-positive and negative infection. Finally, we demonstrate analytically and numerically that HBV RNA is more sensitive than HBV DNA to increases in CAM effectiveness.

**Author summary:** Capsid assembly modulators (CAMs) are a novel class of anti-hepatitis B virus (HBV) treatments in clinical trials. These CAMs have a distinct mechanism of action from nucleos(t)ide analogues and thus represent an attractive option for the treatment of chronic HBV infection. We developed a multiscale model of the intracellular HBV lifecycle and extracellular dynamics using a time-since-infection structured partial differential equation. We fit the model to participant data from a recent phase I trial, performed a detailed parameter sensitivity analysis, identified key mechanisms driving viral response to first-generation CAM treatment, and demonstrated that HBV RNA is more sensitive than HBV DNA to changes in CAM efficacy, highlighting the potential role of HBV RNA as a biomarker for CAM effectiveness.

## Introduction

Despite the availability of an effective vaccine, chronic hepatitis B virus infection (CHB) imposes a major burden on health systems worldwide and is estimated to contribute to one million deaths per year [1]. Often referred to as a silent epidemic [2], the World Health Organization estimated that over 296 million individuals worldwide were living with CHB in 2019 [3]. While effective antiviral therapies, such as pegylated interferon-*α* and nucleos(t)ide analogues (NAs) exist, interferon-*α* treatment is associated with an unfavourable toxicity profile and NA treatment has a low functional cure rate [4, 5]. Specifically, while NA treatment often leads to undetectable hepatitis B viral loads, viral rebound occurs rapidly upon treatment interruption. This viral rebound is driven by the presence of covalently closed circular DNA (cccDNA) that acts as a stable transcriptional template for hepatitis B virus (HBV) replication in infected hepatocytes [6], and thus necessitates life-long treatment. There is consequently a pressing need for the development of novel anti-HBV therapies.

A novel class of HBV antivirals with a distinct mechanism of action from NAs, capsid assembly modulators (CAMs), have demonstrated promising results in recent clinical trials [4, 7–9]. CAMs interfere with a crucial step in the HBV viral life cycle by inhibiting the encapsidation of pregenomic RNA (pgRNA) [4, 9, 10]. By blocking the encapsidation of pgRNA and the resulting production of HBV DNA, CAM treatment has been shown to drive significant declines in HBV RNA and HBV DNA serum concentrations [4, 7, 9, 11]. Here, we consider a phase I trial of the first-generation CAM vebicorvir [4] and we analyse the antiviral efficacy of vebicorvir by developing a multiscale mathematical model of CAM treatment in the context of CHB.

Mathematical modeling has provided extensive insight into the viral dynamics of both hepatitis B and C [12–19]. The majority of existing models focus on extracellular quantities, such as HBV DNA or HBV RNA, which can be immediately compared against clinical data [13]. These models have provided valuable insight into the development of drug-resistance and treatment efficacy in hepatitis C infection [12, 20, 21]. Further, multiscale models, which characterise both the intracelluar and extracelluar viral dynamics, and thus permit a more precise representation of the mechanism of action of novel therapies, have been established to understand viral kinetics following treatment in hepatitis C [22–25]. However, much of the existing modeling in hepatitis B has focused on the dynamics of HBV DNA without explicitly considering the intracellular processes that comprise the HBV viral life cycle [26–32]. This modeling has identified increased death rates of infected hepatocytes in HBe antigen (HBeAg) negative infections, highlighted the role of HBeAg status as a significant predictor of extracelluar viral dynamics, and has characterised the decay kinetics of HBV DNA during treatment. Nevertheless, recent experimental and modeling work has highlighted the role of HBV RNA as an important biomarker in understanding CHB treatment efficacy [33–35]. For example, Goncalves et al. [36] developed a multiscale model of HBV infection that explicitly includes intracellular pgRNA and relaxed circular DNA (rcDNA) dynamics as well as circulating HBV DNA and RNA. They used the model to understand clinical data following treatment with the CAM, RG7907, or the NA, entecavir [36].

Here, we develop a multiscale model of HBV infection similar to the model developed by Goncalves et al. [36] and applied to the CAM RG7907. Specifically, we explicitly consider the dynamics of intracelluar HBV encapsidated pgRNA and rcDNA and tie these dynamics to the extracellular dynamics of HBV RNA and HBV DNA. Unlike Goncalves et al. [36], we incorporate the dynamics of uninfected hepatocytes and alanine aminotransferase (ALT). As shown by [37, 38], ALT dynamics can facilitate parameter identification in mathematical models of HCV infection and is commonly used as a biomarker of liver damage [39, 40]. We fit our multiscale model to the HBV RNA, HBV DNA, and ALT of 29 individuals with CHB who participated in the 28 day, multiple ascending dose, monotherapy trial of vebicorvir [4]. We use our multiscale model to identify the effect of vebicorvir treatment on HBV RNA and HBV DNA dynamics, identify mechanistic differences between HBeAg-positive and HBeAg-negative infections, identify the intracellular mechanisms that contribute to viral decline during treatment, and evaluate HBV RNA and HBV DNA as biomarkers of CAM efficacy.

## Methods

### Viral load data

Our study uses longitudinal viral measurements made on days 0, 1, 7, 14, 21, 28, 35, 42, and 56 from the phase 1, randomized, placebo-controlled, multiple ascending dose study of the first-generation CAM, vebicorvir (NIH Trial identifier: NCT02908191) [4]. Briefly, 32 participants with CHB and no previous HBV therapy in the 3 months preceding the trial received either 100 mg (*n* = 10), 200 mg (*n* = 10), 300 mg (*n* = 10), or 400 mg (*n* = 2) oral doses of vebicorvir daily for 28 days and then followed for another 28 days off therapy. The majority (*n* = 17) of participants were HBeAg-positive with further inclusion criteria reported by Yuen et al. [4].

One of the two participants in the 400 mg dose cohort discontinued treatment following an adverse event [4]. We therefore excluded the 400 mg dose cohort as only one participant completed the trial. In addition, we excluded an individual in the 300 mg dose cohort due to a pre-exisiting known CAM resistance mutation (Thr109Met) [4]. The remaining 29 participants in our study were in the 100 mg (*n* = 10), 200 mg (*n* = 10), and 300 mg (*n* = 9) dosing groups, with six, five, and six HBeAg-positive individuals in the 100 mg, 200 mg, and 300 mg dose cohorts, respectively. We show representative serum HBV RNA and DNA dynamics in Fig. 1.

**Fig 1.**
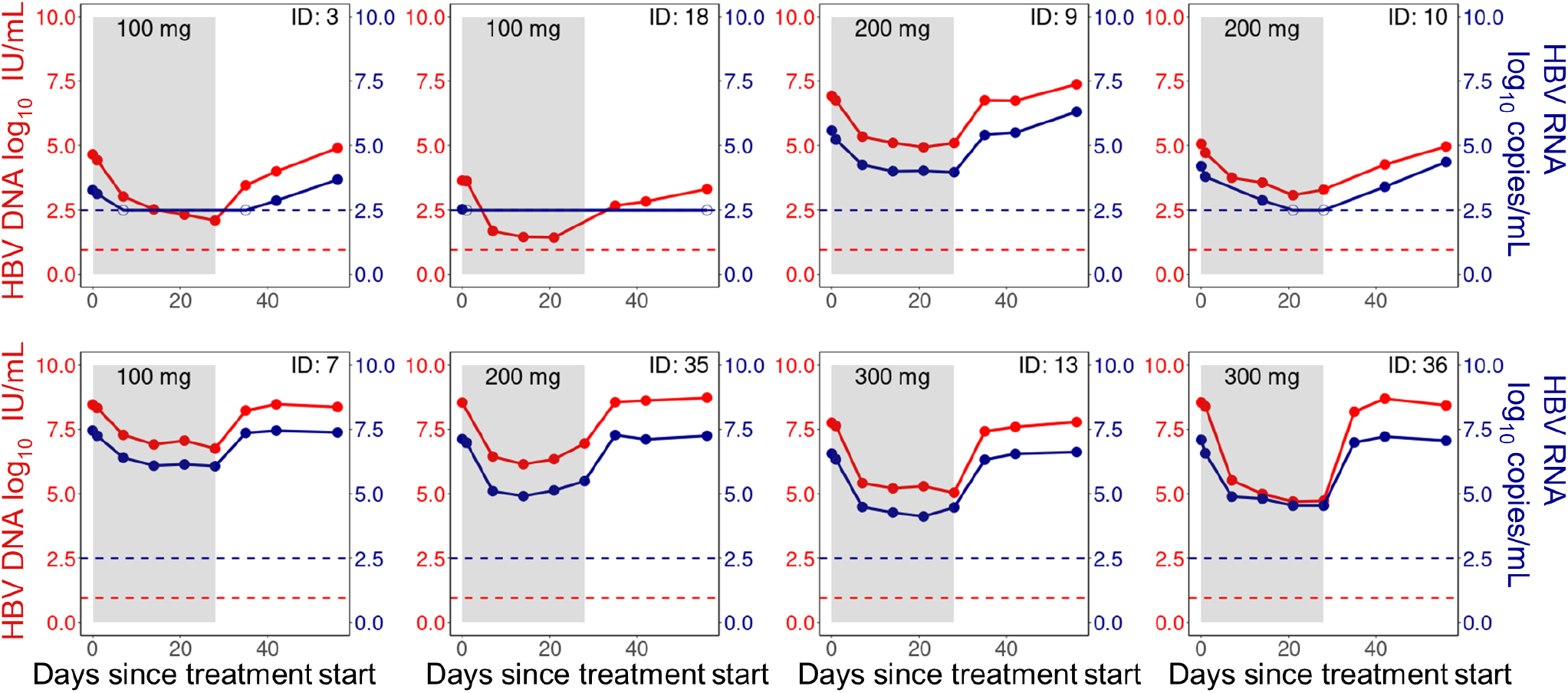
Viral measurements during treatment and follow-up. Longitudinal HBV DNA (red) and HBV RNA (blue) measurements during treatment (shaded region) and follow-up (unshaded region) for selected individuals. Top panel: HBeAg-negative participants, and bottom panel: HBeAg-positive participants. Dots are viral measurements, while solid lines are used to connect the measurements. Horizontal dashed lines represent the lower limit of detection (LLoD) of 0.95 log_10_ IU/mL for HBV DNA (red) and 2.49 log_10_ copies/mL for HBV RNA (blue). Open circles are viral measurements below the LLoD. Individuals were selected to show the heterogeneity in the viral data across both HBeAg-positive and HBeAg-negative individuals.

Vebicorvir treatment decreased serum HBV RNA and HBV DNA concentrations. The mean decrease in HBV RNA and HBV DNA was 1.64 log_10_ copies/mL and 2.12 log_10_ IU/mL, respectively, during the 28 day treatment period. Rebound to approximately pre-treatment baseline serum HBV RNA and HBV DNA concentrations occurred rapidly following treatment cessation. The lower limit of detection (LLoD) for HBV DNA was 0.95 log_10_ IU/mL while the lower limit of quantitation (LLoQ) was 1.28log_10_ IU/mL [4]. To convert from IU/mL to copies/mL, we used the standard conversion factor of 5.82 copies/IU HBV DNA [41]. Finally, Yuen et al. [4] used two independent assays for HBV RNA concentrations; we use the LLoD of 2.49log_10_ copies/mL, which corresponds to the more sensitive of the two assays.

### Multiscale model of chronic HBV infection

Our multiscale model of CHB incorporates the major features of both the intracellular life cycle and extracellular dynamics of HBV. Broadly speaking, we model the intracellular dynamics of encapsidated pgRNA and rcDNA within infected hepatocytes, which allows us to accurately represent the mechanism of action of vebicorvir. Further, we model the extracellular dynamics of infected and uninfected hepatocytes, HBV RNA and DNA, and ALT.

As has been shown previously in hepatitis C [37], modeling ALT dynamics can inform estimates of infected hepatocyte lifespans. Moreover, while hepatocytes are able to proliferate to counter liver damage, we do not expect the pool of hepatocytes to be constant. Therefore, we explicitly include the dynamics of the uninfected hepatocytes in our model. A schematic representation of our model is given in Fig. 2.

**Fig 2.**
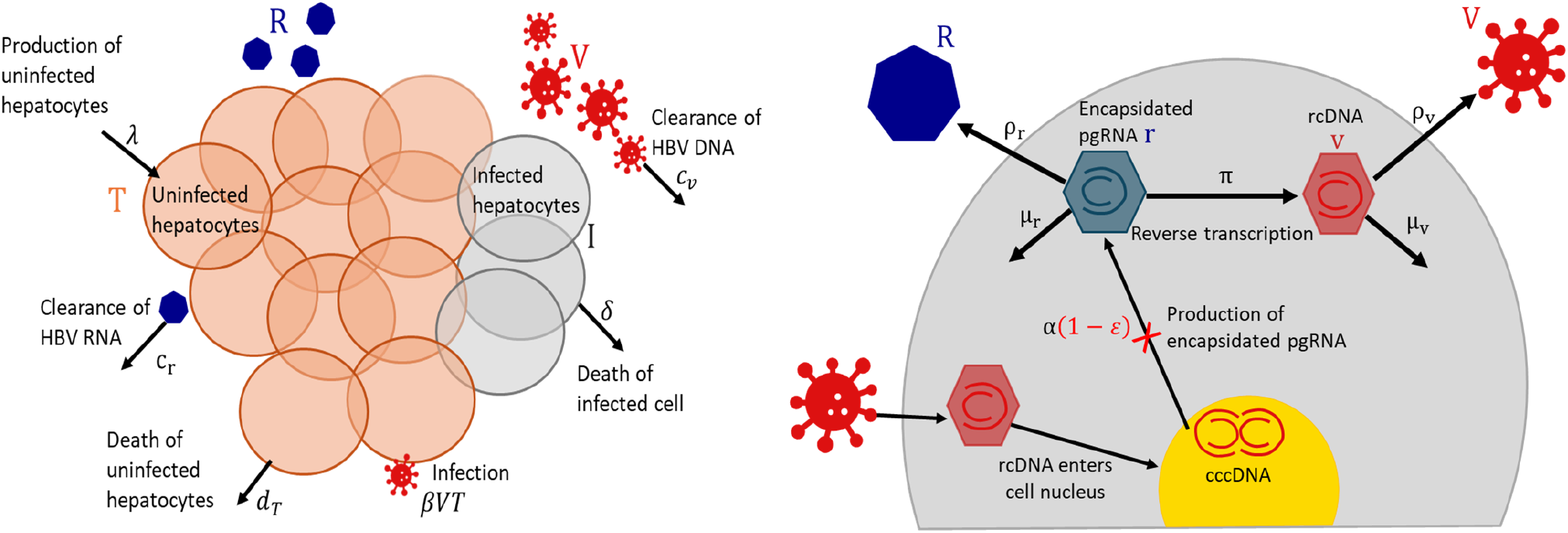
Schematic illustration of the multiscale model. Left panel: HBV extracellular dynamics, where uninfected hepatocytes (*T*) are produced at a constant rate *λ* and die at per capita rate *d*_*T*_. Hepatocytes become infected cells (*I*), following infection by HBV DNA (*V*). Infected cells are lost at per capita rate *δ* and secrete both HBV RNA (*R*) and HBV DNA at constant rates. The HBV RNA and HBV DNA are cleared at rates *c*_*r*_ and *c*_*v*_, respectively. Right panel: HBV intracellular life cycle, which begins with a hepatocyte being infected and the release of rcDNA into the cell cytoplasm following the disintegration of its capsid. This rcDNA enters the cell nucleus and is converted to cccDNA. Encapsidated pgRNA (*r*) is produced by cccDNA at rate *α*. The encapsidated pgRNA is assembled into membrane bound particles and secreted as HBV RNA by the infected cell at rate *ρ*_*r*_, decays at rate *μ*_*r*_, or is reverse transcribed into encapsidated rcDNA (*v*) at rate *π*. The rcDNA is either assembled into viral particles and secreted into the circulation at rate *ρ*_*v*_ or decays at rate *μ*_*v*_ in the cell. Treatment with vebicorvir inhibits the production of encaptidated pgRNA with an effectiveness of *ε* (red cross in the right panel).

At the extracellular scale, uninfected hepatocytes (*T*) are produced at a constant rate *λ* and cleared linearly with per capita rate *d*_*T*_. Hepatocytes are infected by HBV DNA containing particles (*V*) with rate constant *β*. These infected hepatocytes die with per capita death rate *δ* and produce HBV RNA containing particles (*R*) and HBV DNA containing virions, which are cleared linearly at per capita rates *c*_*r*_ and *c*_*v*_, respectively.

As mentioned, we explicitly model the intracellular processes leading to HBV RNA and HBV DNA production. Specifically, we keep track of the time since infection (or infection age) of HBV infected hepatocytes using an age-structured partial differential equation (PDE). The density of infected cells with infection age *a* at time *t* is given by *i*(*t, a*). These infected cells die at rate *δ*. Following infection, HBV rcDNA is converted to cccDNA in the nucleus of infected hepatocytes. This cccDNA forms a stable template for the production of encapsidated HBV pgRNA. We assume the production of encapsidated pgRNA from cccDNA occurs at a constant rate *α* and denote the amount of intracellular encapsidated pgRNA in an infected hepatocyte with infection age *a* by *r*(*t, a*). Encapsidated pgRNA decays intracellularly at rate *μ*_*r*_, is reverse transcribed into encapsidated rcDNA (*v*(*t, a*)) with rate *π*, or is secreted by infected cells as enveloped HBV RNA containing particles into the circulation at the rate *ρ*_*r*_. The rate at which encapsidated pgRNA enters the circulation as HBV RNA is *ρ*_*r*_*P*(*t*), where *P*(*t*) is the total amount of pgRNA in infected cells given by

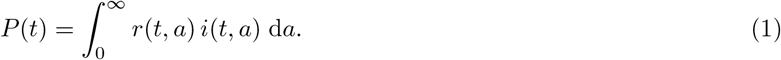

Following reverse transcription of intracellular encapsidated pgRNA, intracellular rcDNA either decays at rate *μ*_*v*_ or is assembled into viral particles and secreted into the circulation as HBV DNA with rate *ρ*_*v*_. The secretion rate of HBV DNA containing particles into the ciruclation is given by *ρ*_*v*_*C*(*t*), where *C*(*t*) represents the total amount of encapsidated rcDNA in infected cells, given by

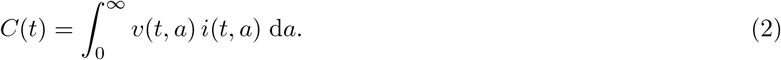

Finally, we explicitly model the dynamics of ALT. Following Ribeiro et al. [38], we assume that ALT is produced at a constant rate *s*, is cleared linearly with rate *c*_*A*_, and is released at a constant rate *α*_*A*_*δ* due to the death of infected cells.

Taken together, the equations describing the multiscale model are

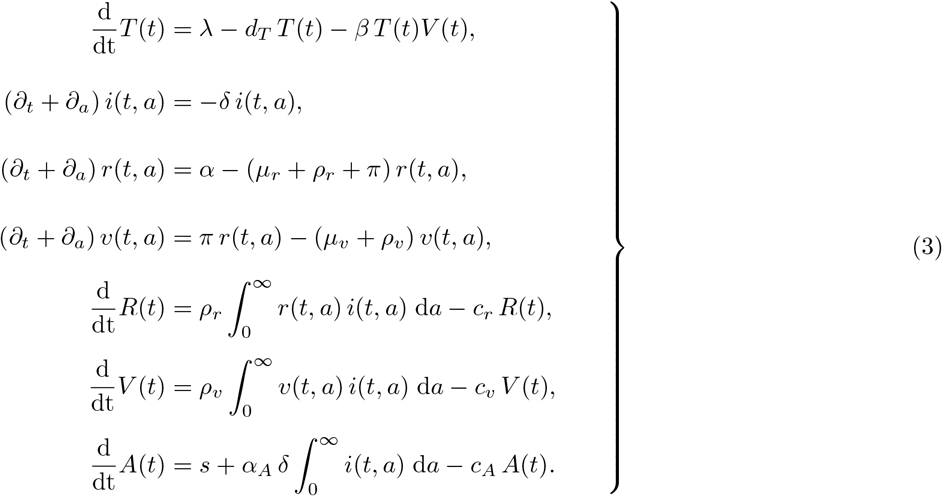

Newly infected cells have infection age *a* = 0, and the density of newly infected cells at time *t* is given by *i*(*t*, 0) = *βV*(*t*)*T*(*t*). We assume that newly infected cells have no intracellular pgRNA or rcDNA, i.e., *r*(*t*, 0) = 0 and *v*(*t*, 0) = 0, as the rcDNA from the initial virion that infected a hepatocyte is transported to the nucleus and converted to cccDNA, and thus behave differently then newly produced rcDNA that can be converted into virions. We discuss the initial conditions of Eq. (3) in the following.

In principle, the rate constants describing intracellular processes within an infected cell as well as the death rate of infected cells could depend on the infection age of the cells. For example, Nelson et al. [42], examined an HIV model in which the rate of viral production and the death rate of productively infected cells varies with their infection age. Similarly, Hailegiorgis et al. [43] developed an agent-based model of acute HBV infection in which the rate of virion production increased until reaching a constant rate. As little information is available on the age-dependence of the parameters in our model, we restrict our analysis to the case of age-independent parameters. In this case, the multiscale PDE model Eq. (3) can be transformed into the following ordinary differential equation (ODE) system using standard techniques [18, 19, 36, 44]

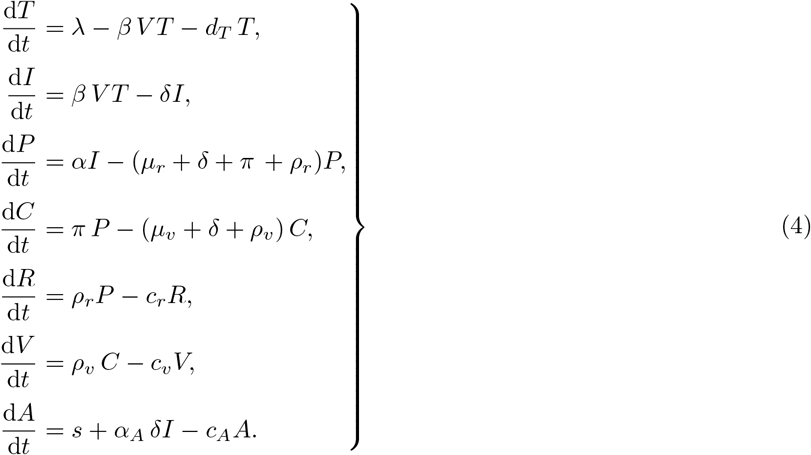

Here, *I*(*t*) is the total concentration of infected hepatocytes defined by

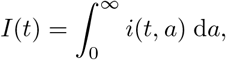

while *P*(*t*) and *C*(*t*) are given in Eq. (1) and (2), respectively. The ODE model Eq. (4) is a mathematically equivalent and numerically tractable representation of the PDE model Eq. (3) under the assumption of age-independent parameters. We refer to the ODE system Eq. (4) as the *pre-treatment/baseline model* throughout this study.

### Modeling vebicorvir pharmacodynamics

In the phase I trial of vebricorir [4], participants received vebicorvir once daily for 28 days. We neglect drug pharmacokinetics during the daily dosing period as the drug was rapidly absorbed [4]. We assume that vebicorvir inhibits pgRNA encapsidation immediately following dosing. Thus, during the treatment period, we model the pharmacodynamic effects of vebicorvir by inhibiting the production rate of encapsidated pgRNA, *α*, by the factor (1 − *ε*(*t*)), where *ε*(*t*) ∈ [0, 1] represents the CAM effectiveness at time *t*.

During drug washout, we assume that the vebicorvir concentration decays exponentially at the rate *k* from a dose-dependent steady-state concentration *C*^∗^. Then, during drug washout, we use a maximum effect, or Emax model, for vebicorvir effectiveness given by

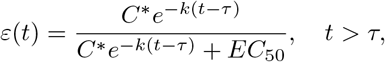

where *τ* = 28 days is the duration of the treatment period. Setting *ε*(*τ*) = *ε*_*c*_, where *ε*_*c*_ is the drug effectiveness during therapy, gives

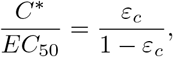

so

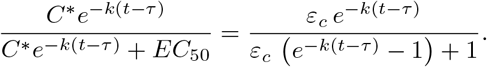

Incorporating the waning vebicorvir effects during drug wash-out only necessitates estimating the clearance rate, *k*. We also considered an approach where the drug efficacy, *ε*(*t*), is set to zero immediately after treatment cessation, and a scenario where *ε*(*t*) decays exponentially for *t > τ*.

Incorporating vebicorvir mediated inhibition of pgRNA encapsidation into the baseline model Eq. (4) gives

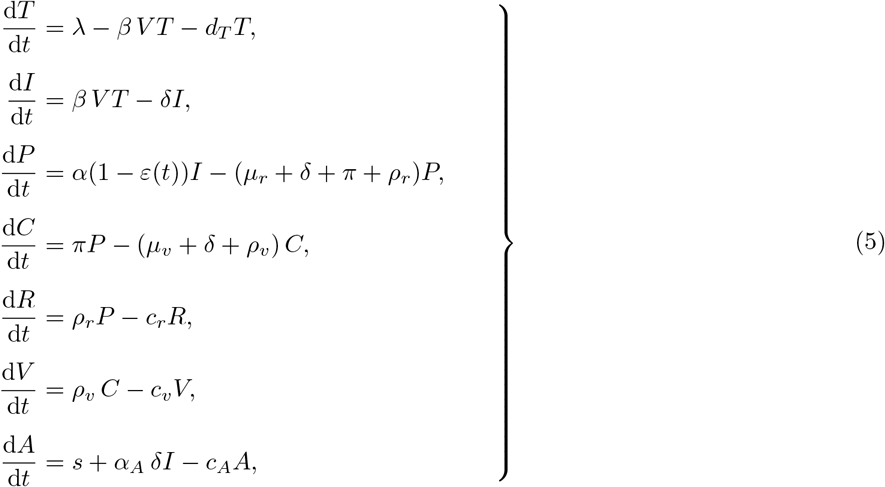

where the drug efficacy, *ε*(*t*), is defined by

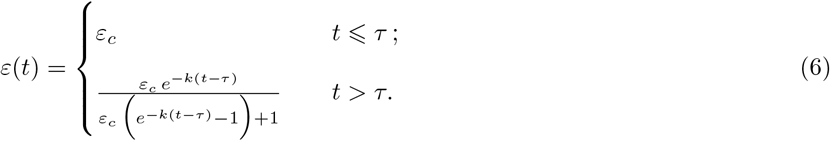

We refer to the ODE model Eq. (5) as the *treatment model*. The model parameters, units, and biological descriptions are summarised in Table 1.

**Table 1.** Model parameters and descriptions.

| Parameter (unit) | Description |
| --- | --- |
| $\lambda$ (cells/mL/day) | Production rate of hepatocytes |
| $d_T$ (/day) | Death rate of uninfected hepatocytes |
| $\beta$ (mL/copies/day) | Infection rate constant |
| $\varepsilon(t)$ | Drug effectiveness in preventing pgRNA encapsidation |
| $\delta$ (/day) | Death rate of infected cells |
| $\alpha$ (copies/cell/day) | Production rate of encapsidated pgRNA |
| $\pi$ (/day) | Reverse transcription rate of pgRNA to rcDNA |
| $\rho_r$ (/day) | Secretion rate of intracellular encapsidated pgRNA as HBV RNA |
| $\rho_v$ (/day) | Viral assembly/secretion rate |
| $\mu_r$ (/day) | Intracellular decay rate of encapsidated pgRNA |
| $\mu_v$ (/day) | Intracellular decay rate of rcDNA |
| $c_r$ (/day) | Extracellular clearance rate of HBV RNA |
| $c_v$ (/day) | Extracellular clearance rate of HBV DNA |
| $s$ (U/L/day) | Baseline production rate of ALT |
| $\alpha_A$ (U/L/mL/cell) | Amount of ALT released by infected cells upon their death |
| $c_A$ (/day) | Clearance rate of ALT |
| $\varepsilon_c$ | Drug effectiveness during treatment |
| $k$ (/day) | CAM clearance rate |
| $\tau$ (days) | Treatment duration |

### Initial conditions corresponding to chronic HBV infection

Since we are interested in chronic HBV infection, we assume that the viral dynamics model is in steady-state prior to treatment. We thus use the steady-state solutions of the baseline model Eq. (4) as the initial conditions of the treatment model Eq. (5). The baseline viral load, *V*_0_, is given by

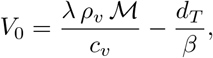

where ℳ = *π α/*(*δ ψ*_1_ *ψ*_2_), with *ψ*_1_ = *μ*_*r*_ + *δ* + *π* + *ρ*_*r*_ and *ψ*_2_ = *μ*_*v*_ + *δ* + *ρ*_*v*_. The remaining steady-state concentrations can be written in terms of *V*_0_ as follows

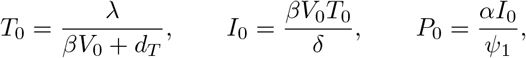

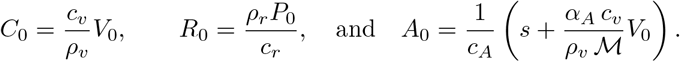

We calculate these expressions explicitly in terms of the model parameters in S1 Text. We consider *t* = 0 as the beginning of the clinical trial and set

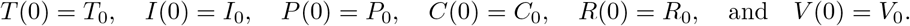

Finally, we note that imposing that the viral dynamics model is in steady-state prior to treatment yields natural candidates for initial densities *i*_0_(*a*), *r*_0_(*a*), and *v*_0_(*a*) of the age-structured PDE model. Specifically, the initial density of infected hepatocytes at time *t* = 0 represent precisely those hepatocytes that were infected at *t <* 0 and have not been cleared in the intervening time. Following Cassidy et al. [45], it is possible to map the initial densities backwards along the characteristic lines and, using the assumption of chronic infection, obtain explicit expressions for *i*_0_(*a*), *r*_0_(*a*), and *v*_0_(*a*) as functions of the baseline viral load and uninfected hepatocyte concentration.

### Data fitting and parameter estimation

We fit the multiscale model (Eq. (5)) to the longitudinal HBV RNA, HBV DNA and ALT measurements of the 29 participants from Yuen et al. [4] in our study using a population-based nonlinear mixed-effects modeling framework implemented in Monolix 2021R1 [46]. As mentioned, we convert the HBV DNA output from our model to IU/mL using the conversation rate of 5.82 copies to 1 IU when fitting the model to the data [41].

Parameter estimation was performed by maximizing the likelihood estimator using the stochastic approximation expectation-maximization (SAEM) algorithm [47] implemented in Monolix software 2021R1 [46]. The log-likelihood was calculated using the importance sampling Monte Carlo method. HBV RNA measurements below the LLoD of 2.49log_10_ copies/mL and HBV DNA measurements below the LLoD of 0.95log_10_ IU/mL were left-censored.

### Fixed parameters

We fixed some of the model parameters to estimates from the literature to reduce the number of free parameters in our model. Based on the estimate that, in the absence of infection, the liver has 2 × 10^11^ hepatocytes [48], we assumed the hepatocyte concentration is *T*_*ue*_ ≈ 1.3 × 10^7^ cells/mL, as was previously done by Neumann et al. [12].

Uninfected hepatocytes have a roughly 6 month (∼ 180 days) half-life [49], which corresponds to a death rate of *d*_*T*_ = log(2)*/*180 ≈ 0.004 /day. In the absence of infection, the steady-state concentration of hepatocytes is given by *T*_*ue*_ = *λ/d*_*T*_ so *λ* = *d*_*T*_ *T*_*ue*_ = 0.004 × (1.3 × 10^7^) cells/day.

Biologically, one would expect that the clearance rates of HBV RNA (*c*_*v*_) particles and of HBV DNA (*c*_*r*_) particles are similar, so we assume *c*_*v*_ = *c*_*r*_ = *c* (we note that this was also used by Goncalves et al. [36]) Estimating the clearance rate *c* is challenging from our clinical data, so we follow Goncalves et al. [36] and test five different fixed values of *c* = 1, 2, 5, 15, and 20 /day based on previous studies [29, 50]. We selected the value of *c* that minimizes the corrected Bayesian information criterion (BICc) when fitting the viral dynamics model to the clinical data [51].

Similarly, as in Goncalves et al. [36], we assumed that encapsidated pgRNA and rcDNA are degraded at identical rates in infected cells, so *μ*_*v*_ = *μ*_*r*_ = *μ*. As mentioned by Goncalves et al. [36], the intracellular degradation rate is difficult to estimate from circulation data alone. We thus tested 11 fixed values of *μ* = 0, 0.1, 0.2, …, and 1.0 /day and used the value of *μ* that gave the lowest negative log-likelihood.

### Estimated parameters

At the extracellular scale, we estimated the infection rate (*β*) and the death rate of infected cells (*δ*). We also estimated the intracellular production rate of encapsidated pgRNA (*α*), the reverse transcription rate of pgRNA to rcDNA (*π*), and the secretion rates of HBV RNA (*ρ*_*r*_) and HBV DNA (*ρ*_*v*_) by fitting Eq. (5) to the HBV RNA, HBV DNA, and ALT concentrations. Finally, we also estimated both the effectiveness, *ε*, and clearance rate, *k*, of vebicorvir.

We also fit the pre-treatment baseline level of ALT (*A*_0_), its clearance rate (*c*_*A*_), and the uninfected steady-state ALT concentration (*A*_*ue*_). To estimate the baseline production rate of ALT (*s*), we consider the ODE for ALT in Eq. (5) in the absence of infection

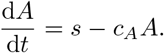

Thus, we calculate the baseline production rate *s* = *c*_*A*_ *A*_*ue*_. Then, at the pre-treatment steady-state, the amount of ALT released from dead or dying infected cells is given by

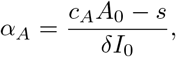

where *I*_0_ is the baseline number of infected hepatocytes at treatment initiation and *A*_0_ is the ALT concentration at treatment initiation, which we estimated from data. We give the details of our structural, error, and covariate model in the S1 Text.

### Parameter sensitivity analysis

We performed a local sensitivity analysis for each participant by varying estimated parameters by ±10% from the fit values. We measured the resulting change in the viral nadir and the time to viral rebound, where we defined viral rebound as the first time that HBV DNA concentrations reach 85% of pre-treatment viral load. To translate these individual sensitivity analyses, we considered the median change over all participants. Finally, we adapted the continuation technique from [52] to quantify the dependence of each estimated parameter on the viral load data.

## Results

### Model fits to participant data

We fit our mathematical model (Eq. (5)) to the dynamics of HBV RNA, HBV DNA and ALT simultaneously and show the individual fits of our model to the HBV RNA and DNA data in Figs. 3 and 4. The corresponding individual fits to ALT are shown in Figs. L and M of S1 Text. Here, we emphasize that we simultaneously fit the data from all participants in the Yuen et al. [4] trial using a nonlinear mixed effects framework which simultaneously considers the totality of the data. In general, our model captures the viral dynamics in all participants both during treatment and during the viral rebound that follows treatment cessation.

**Fig 3.**
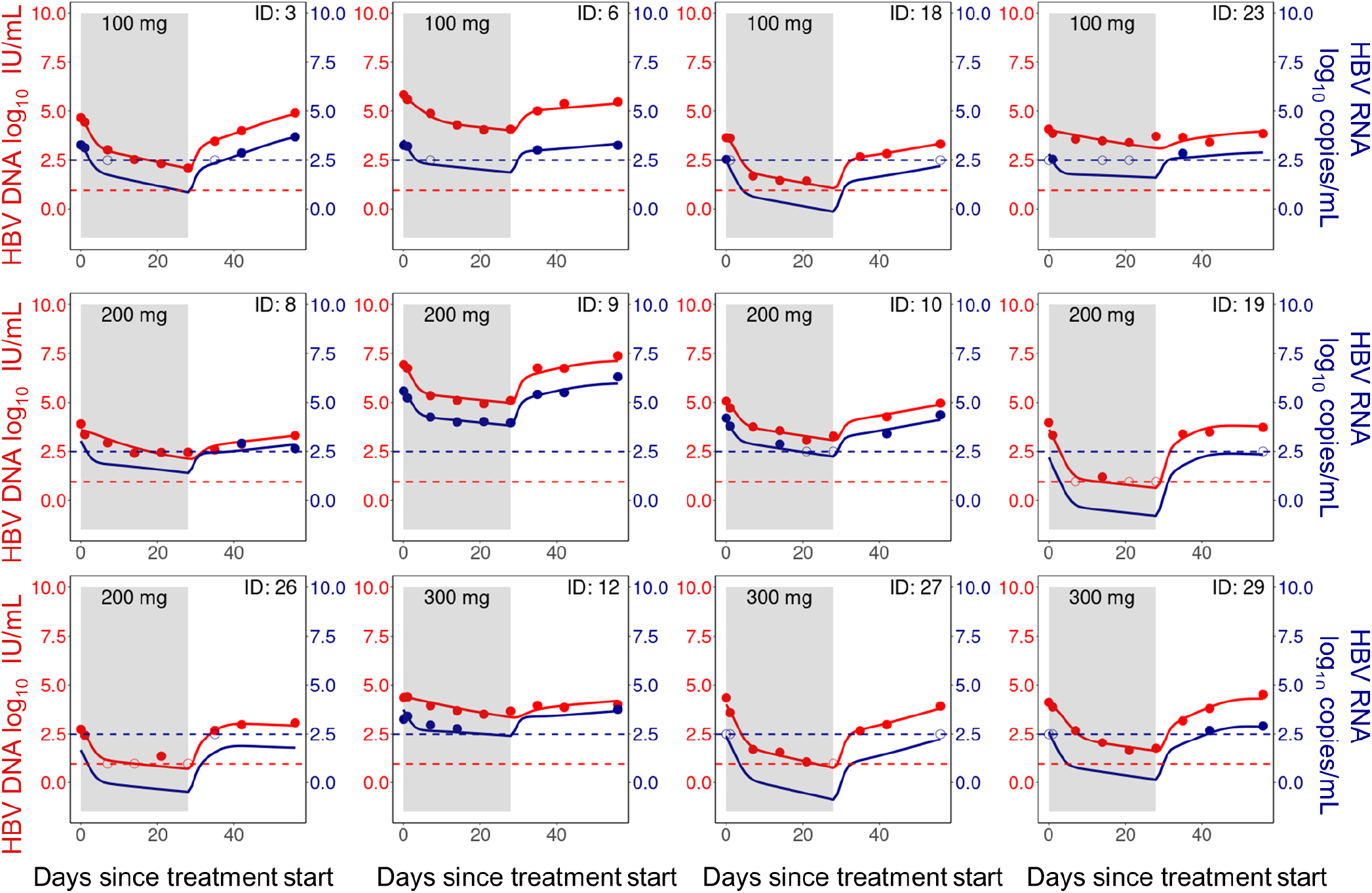
Model fits of HBV RNA and HBV DNA (HBeAg-negative group). Individual fits to the longitudinal HBV RNA (blue) and HBV DNA (red) measurements during treatment (shaded area) and follow-up. Dots are viral measurements, and solid lines are model predictions. Horizontal dashed lines represent the lower limit of detection (LLoD) of 0.95 log_10_ IU/mL for HBV DNA (red) and 2.49 log_10_ copies/mL for HBV RNA (blue). Open circles are viral measurements below the LLoD. The corresponding ALT fits for these participants are given in S1 Text Fig. H.

**Fig 4.**
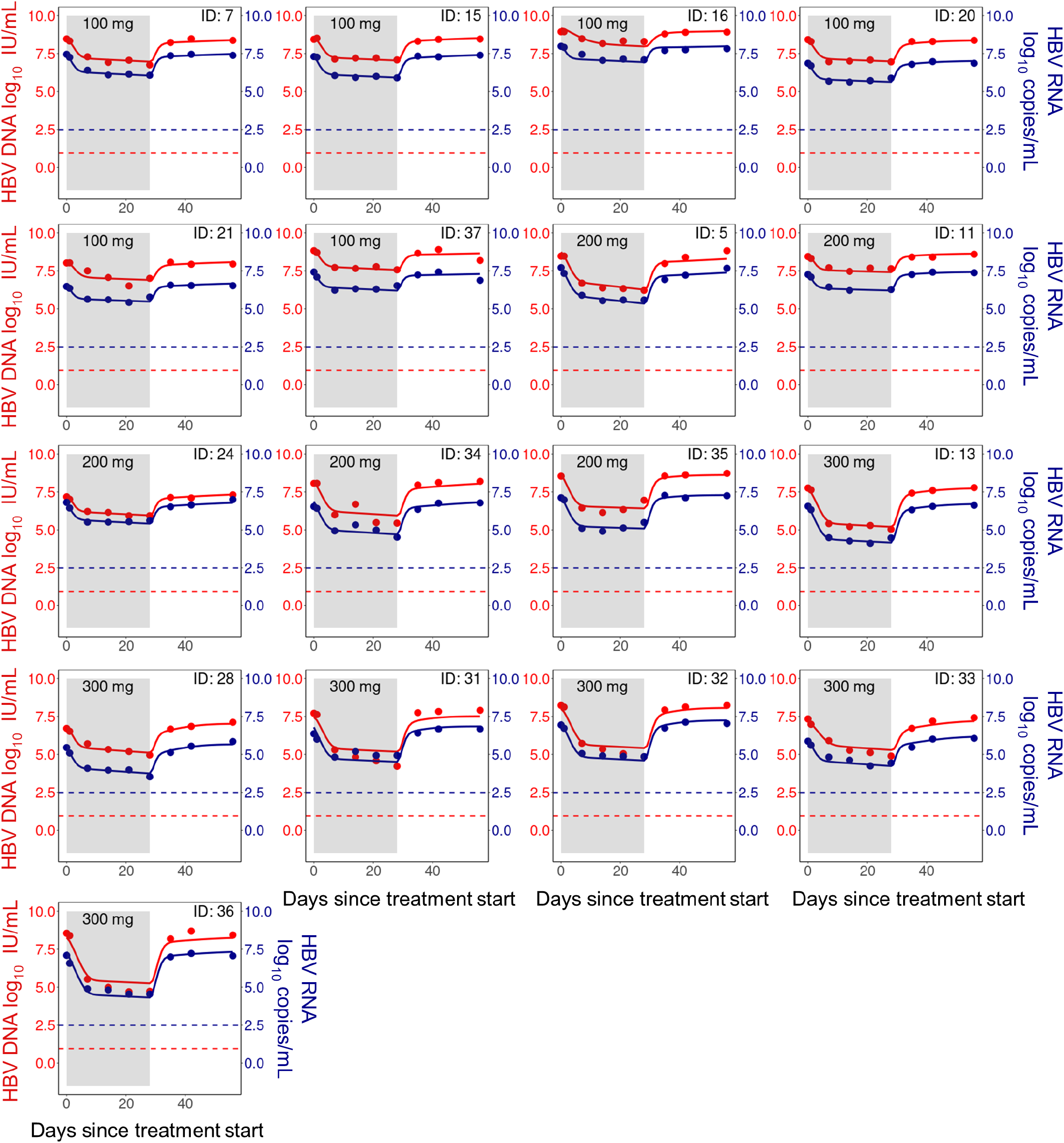
Model fits of HBV RNA and HBV DNA (HBeAg-positive group). Individual fits to the longitudinal HBV RNA (blue) and HBV DNA (red) measurements during treatment (shaded area) and follow-up. Dots are viral measurements, and solid lines are model predictions. Horizontal dashed lines represent the lower limit of detection (LLoD) of 0.95 log_10_ IU/mL for HBV DNA (red) and 2.49 log_10_ copies/mL for HBV RNA (blue). Open circles are viral measurements below the LLoD. The corresponding ALT fits for these participants are given in Fig. I of S1 Text.

The population-level parameters were well-estimated, as measured by relative standard error, and reported in Table 2. The best model fits, as measured by BICc, were obtained for *c*_*r*_ = *c*_*v*_ = 1/day. This estimated clearance rate is much smaller than the estimate reported by Goncalves et al. [36] using a similar model for a phase I trial of the CAM RG7907. There, Goncalves et al. [36] reported *c* = 20/day, with similar model fits obtained for *c* = 5/day and *c* = 10/day. However, the rapid decay of HBV RNA and HBV DNA predicted by large values of *c* ⩾ 5/day is incompatible with our viral dynamics data from days 1 and 2 post-treatment initiation. Indeed, our fitting and exploration of parameter space indicated a strong preference for *c* ⩽ 5/day. We note that the viral dynamics data considered by Goncalves et al. [36] did not include HBV RNA measurements taken before day 7 post-treatment initiation, which may explain the differences in our estimates of *c*. We also identified a dose-dependent vebicorvir effect, with estimated efficacies of 91.9%, 96.3%, and 98.8% for the 100, 200, and 300 mg daily dose. Finally, the value for the degradation rate of intracellular pgRNA and rcDNA that led to the lowest value the log-likelihood was *μ* = 0.

**Table 2.** Estimated population parameters. Random effects is specified as NA where a parameter is fixed or estimated at the population level. P-values were computed using the Wald test in Monolix and used to compare the population estimates for covariates.

| Parameter | Fixed Effects (R.S.E., %) | Random Effects (R.S.E., %) |
| --- | --- | --- |
| $\varepsilon_c$ (100 mg) | 0.919 (1.57) | 2.06 (22.5) |
| $\varepsilon_c$ (200 mg)<sup>*1</sup> | 0.963 (51.5) | - |
| $\varepsilon_c$ (300 mg)<sup>*2</sup> | 0.988 (33.1) | - |
| $\beta$ (HBeAg-negative)<sup>**</sup> | $6.8 \times 10^{-7}$ mL/copies/day (6.08) | 1.14 (22.6) |
| $\beta$ (HBeAg-positive) | $9.9 \times 10^{-11}$ mL/copies/day (13.4) | - |
| $\alpha$ (HBeAg-negative)<sup>***</sup> | 0.203 copies/cell/day (36.6) | 0.84 (21.3) |
| $\alpha$ (HBeAg-positive) | 365.6 copies/cell/day (10.2) | - |
| $\pi$ | 205.6 /day (26.8) | 0.32 (35.1) |
| $\delta$ (HBeAg-negative)<sup>†</sup> | 0.062 /day (16.8) | 0.39 (23.2) |
| $\delta$ (HBeAg-positive) | 0.022 /day (20.1) | - |
| $\rho_v$ | 1.87 /day (106) | 1.65 (57.3) |
| $\rho_r$ | 2.37 /day (145) | 0.33 (NaN) |
| $c$ | 1 /day (fixed) | NA |
| $\mu$ | 0 /day (fixed) | NA |
| $c_A$ | 0.067 /day (17.6) | NA |
| $A_0$ | 38.4 U/L (3.47) | 0.29 (13.3) |
| $A_{ue}$ | 23.2 U/L (5.52) | 0.29 (23.5) |
| $k$ | 1.69 /day (27.0) | 0.20 (115) |

<sup>\*1</sup> $p = 0.052$ , <sup>\*2</sup> $p = 0.0025$ , <sup>\*\*</sup> $p = 7.3 \times 10^{-14}$ , <sup>\*\*\*</sup> $p < 2.2 \times 10^{-16}$ , <sup>†</sup> $p = 6.2 \times 10^{-7}$ .

We tested if the parameters determining intracellular dynamics are well-informed by the extracellular measurements. We first characterised the sensitivity of two clincally relevant measures, the viral nadir and time to viral rebound, to small perturbations in the parameters representing intracellular mechanisms and show the results of our local sensitivity analysis in Fig. F. We found that these two measurements are sensitive to the parameters determining intracellular dynamics. Unsurprisingly, the viral nadir was sensitive to the production rate of encapsidated pgRNA, *α*, CAM effectiveness, *ε*, and the death rate of infected hepatocytes, *δ*. However, the viral nadir was insensitive to the infection rate *β*, which indicates that viral dynamics during treatment being driven by hepatocytes that were infected prior to treatment initiation. As virus production rapidly resumes following treatment cessation, the time to viral rebound was relatively insensitive to the CAM effectiveness, *ε*. We also found that an increased production rate of encapsidated pgRNA, *α*, decreased the time to viral rebound as would be expected. Conversely, increasing the death rate of infected hepatocytes, *δ*, leads to fewer infected hepatocytes at treatment cessation and thus prolonged the time to rebound. Both the viral nadir and time to rebound were sensitive to the clearance rate *c* = *c*_*v*_ = *c*_*r*_. Finally, as the estimates for *δ* and *α* strongly influence the viral nadir and time to rebound, we conclude that the parameters *δ, α, c* and *ε* are informed by the viral dynamics during therapy.

However, the viral nadir and time to rebound are insensitive to the reverse transcription rate, *π*, and secretion rates, *ρ*_*v*_ and *ρ*_*r*_. We therefore next tested if small perturbations in the observed data will influence individual parameter estimation using the likelihood continuation method [52]. We found that the estimates of the export rates *ρ*_*r*_ and *ρ*_*v*_ are sensitive to 10% changes in each of the HBV RNA and HBV DNA measurements. These parameters are particularly sensitive to measurements during the final two weeks of treatment (Fig. G of S1 Text). Furthermore, these later measurements also inform the estimate of the reverse transcription rate, *π*. The likelihood continuation analysis indicates that these model parameters are therefore informed by the measurements during treatment [52]. Lastly, the initial conditions denoting chronic HBV infection are directly informed by the baseline viral load and strongly depend on *β*. Taken together, our analysis indicates that our parameter estimates for the intracellular dynamics are well-informed by the available extracellular data.

We found no significant difference in the baseline ALT levels between the HBeAg-positive and HBeAg-negative groups, even though two HBeAg-positive individuals (ID: 5, 24) had elevated ALT at the start of treatment. The elevated baseline ALT levels in these individuals may signify an adaptive anti-HBV immune response. Their ALT levels declined during treatment and approached a similar level to the other participants by day 56 post-treatment initiation (Fig. I of S1 Text). There were no significant changes in the ALT levels of the remaining participants throughout the study period.

### Mechanistic differences between HBeAg-positive and negative infection

HBeAg status is an important predictor of clinical progression with faster progression to liver disease observed in HBeAg-negative patients, despite persistently higher viral load in HBeAg-positive patients [29, 53]. We leveraged our multiscale model to identify the mechanistic differences between HBeAg-positive and HBeAg-negative participants. As expected, the baseline HBV DNA concentration, *V*_0_, is significantly higher in HBeAg-positive participants (1.4 × 10^8^ vs 2.3 × 10^4^ IU/mL, *p* = 7.7 × 10^−8^).

We also systematically tested for a covariate effect of HBeAg status on all our model parameters and found that it is a significant covariate on three model parameters, *β, α*, and *δ* (Table 2). Infected hepatocytes typically harbour higher cccDNA concentrations in HBeAg-positive infections [54], which provides a biological mechanism underlying the increased encapsidated pgRNA production rate, *α*. Further, as HBeAg-positive infections are typically linked to immune tolerance, the higher death rate of infected hepatocytes, *δ*, during HBeAg-negative infections may indicate an antiviral immune response [55, 56]. In S1 Text, we calculate the basic reproduction number of our model as

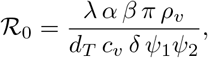

where we recall that *ψ*_1_ = *μ*_*r*_ + *δ* + *π* + *ρ*_*r*_ and *ψ*_2_ = *μ*_*v*_ + *δ* + *ρ*_*v*_.

Although the estimated infection rate (*β*) is lower in the HBeAg-positive group (Table 2), we found that R_0_ is larger for HBeAg-positive participants than for HBeAg-negative participants, with mean estimates of 32.1 (R.S.E = 0.27) and 22.6 (R.S.E. = 0.15) for HBeAg-positive and HBeAg-negative participants, respectively. As the infection rate, *β*, is directly proportional to the basic reproduction number, this result may initially seem counter-intuitive. However, the significantly faster production rate of encapsidated pgRNA, *α*, and the lower death rate of infected cells, *δ*, in HBeAg-positive participants counterbalances the lower infection rate and results in a larger R_0_ estimate for the HBeAg-positive participants.

As mentioned above, HBeAg-positive infections tend to lead to higher viral loads. Our model also predicts increased levels in the mean predicted pre-treatment steady-states for the concentrations of infected hepatocytes, intracellular pgRNA and rcDNA, and HBV RNA and DNA. We show the distribution of individual pre-treatment steady-states in Fig. K of S1 Text.

### Analytical solution of viral dynamics model identifies mechanisms driving biphasic decay

Immediately following treatment initiation, both HBV RNA and HBV DNA concentrations exhibited observable biphasic declines. The rapid first phase of decline is characterized by a half-life of approximately 17 hours for both HBV RNA and HBV DNA and lasts for roughly 7 days, where this half-life is determined by log(2)*/c*. The second, slower phase of decline differed between HBeAg-positive and HBeAg-negative individuals, with estimated half-lives of 32 and 11 days, respectively, determined by log(2)*/δ*.

Under the assumption that HBV DNA concentrations fall sufficiently rapidly during vebicorvir treatment to neglect *de novo* cell infections during treatment, we solved the multiscale model Eq. (5) analytically in the S1 Text. We compare the predicted HBV DNA dynamics obtained by simulating the full ODE model Eq. (5) and the approximation obtained by neglecting new infections for each participant in Figs. A and B of the S1 Text for the HBV DNA and RNA, respectively. The difference between the approximate and full model predictions is less than 0.3 log_10_ for all participants during the 28 day trial. However, the difference between the exact and approximate solutions are generally less than 0.1 log_10_ when the drug effectiveness is high, i.e., for the 200 mg and 300 mg doses.

The analytical solution for HBV RNA concentration after the start of treatment, derived in Eq. (S17) of S1 text, is the sum of three exponentially decaying terms with rates *ψ*_1_, *c*_*r*_, and *δ*, whereas the concentration of HBV DNA is the sum of four exponentially decaying terms with rates *ψ*_1_, *ψ*_2_, *c*_*r*_, and *δ* as given in Eq. (S18) of S1 text. For both HBV RNA and HBV DNA, only two of these phases of decline are observable on the time scale of our study. As mentioned, the first observable phase of both HBV RNA and HBV DNA decline is dominated by clearance of HBV RNA and DNA from the circulation with rate *c*, while the second phase is dominated by the death of infected cells at HBeAg-status dependent rate *δ*. We show the theoretical decay curves in Fig. 5 to illustrate the transition between these two decay phases.

**Fig 5.**
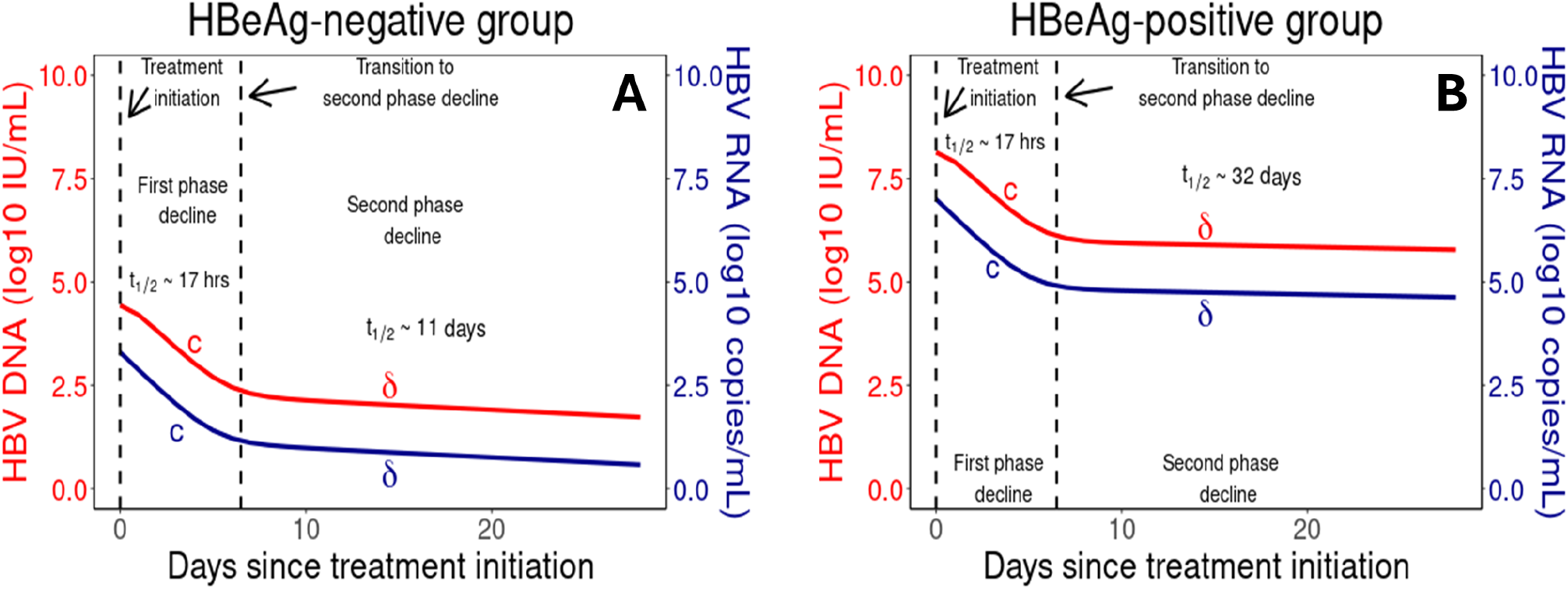
Biphasic decay of HBV RNA and HBV DNA during vebicorvir treatment. Panels A and B show the HBV DNA (red) and HBV RNA (blue) decay profiles during 300 mg daily vebicorvir treatment for the population parameter estimates for HBeAg-negative and positive participants, respectively.

These exponential decays are directly related to the mechanism of action of vebicorvir. As vebicorvir inhibits the encapsidation, and thus production, of encapsidated pgRNA, the intracellular encapsidated pgRNA concentration declines with rate *ψ*_1_ due to degradation, secretion, and reverse transcription, and rapidly reaches a treated quasi-equilibrium 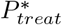 within infected hepatocytes. There is a corresponding decline to a treated quasi-equilibrium 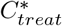 with rate *ψ*_2_ in intracellular rcDNA concentrations. The convergence to these treated quasi-equilibria are sufficiently rapid to be unobservable in the circulating HBV RNA and DNA data.

Then, as 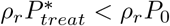 and 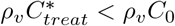, there is a corresponding fall in the secretion of HBV RNA and HBV DNA. Recalling that the system was at steady-state prior to treatment with *ρ*_*v*_*P*_0_ = *cR*_0_ and *ρ*_*v*_*C*_0_ = *cV*_0_, the rapid convergence to the treated quasi-equilibria, 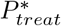 and 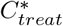, for intracellular pgRNA and rcDNA implies that the HBV RNA and DNA dynamics are initially dominated by clearance, with rate *c*, during the initial phase of decline following treatment initiation. Then, due to the significant decrease of HBV DNA during the first phase of decline, the virus is not able to maintain the infected hepatocyte population at the pre-treatment level via secondary infections. Thus, the death of infected cells drives the second phase of decline and leads to viral decline at the death rate of infected hepatocytes.

The intracellular pgRNA and rcDNA declines to the treated quasi-equilibria has a half-life of log(2)*/ψ*_1_ = 0.003 days, for both HBeAg positive and negative participants, while the intracelluar rcDNA decline has a half-life of log(2)*/ψ*_2_ = 0.37 days and 0.36 days for HBeAg positive and negative participants, respectively. While these declines are too rapid to be observable in circulating HBV RNA and DNA with daily sampling, they could potentially inform an improved understanding of the intracellular HBV life cycle. For example, by measuring the number of rcDNA copies per infected hepatocyte from a pre-treatment liver biopsy and estimating the percentage of infected hepatocytes, we could estimate the baseline total concentration of rcDNA, *C*_0_. Then, assuming that we were able to observe *ψ*_2_, recalling that the first and second phase of HBV RNA and DNA decline directly inform *c* and *δ*, respectively, and that *V*_0_ is typically measured in clinical studies, we find an explicit expression for the intracelluar rcDNA decay rate

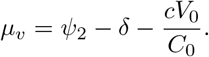

### HBV RNA is more sensitive than HBV DNA to CAM efficacy

Due to the mechanism of action of CAMs in blocking intracellular pgRNA production, HBV RNA dynamics are a potential direct biomarker of target engagement, and thus, drug effectiveness [5, 57]. Here, we use our mathematical model to understand the relationship between CAM efficacy and the observed decay in HBV RNA and HBV DNA.

We performed an analytical sensitivity analysis of the response of HBV RNA and DNA to increase in CAM effectiveness. As previously mentioned, under the assumption that CAM treatment was sufficiently potent to neglect new infections, we solved the multiscale model Eq. (5). In S1 Text, we used this analytical solution to evaluate the impact of parameter changes on model predictions and show that perturbations of CAM effectiveness result in larger relative changes in HBV RNA concentrations than in HBV DNA concentrations. Specifically, we show

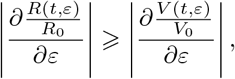

where *R*_0_ and *V*_0_ are the pre-treatment steady-state HBV RNA and HBV DNA levels, respectively, and *ε* is the drug effectiveness. This analytical result supports using HBV RNA dynamics as a biomarker of treatment efficacy, as has been suggested recently [34].

In Fig. 6, we compare the predicted relative changes in log_10_ concentrations of HBV RNA and HBV DNA during the first 14 days of treatment with vebicorvir. We use the mean population parameter estimates for HBeAg-negative and HBeAg-positive participants as the viral dynamics parameters and the population estimates for 100, 200, or 300 mg of vebicorvir for the values of *ε* in Panels A and B, respectively. In all cases, we see that the predicted fold decline in HBV RNA concentration is larger than the corresponding prediction in HBV DNA, which confirms our analytical results. In Fig. E of S1 Text, we show the same comparison for each participant. In all cases, while the decay dynamics of HBV RNA and HBV DNA are similar, HBV RNA undergoes a larger relative decay during the first 14 days of treatment.

**Fig 6.**
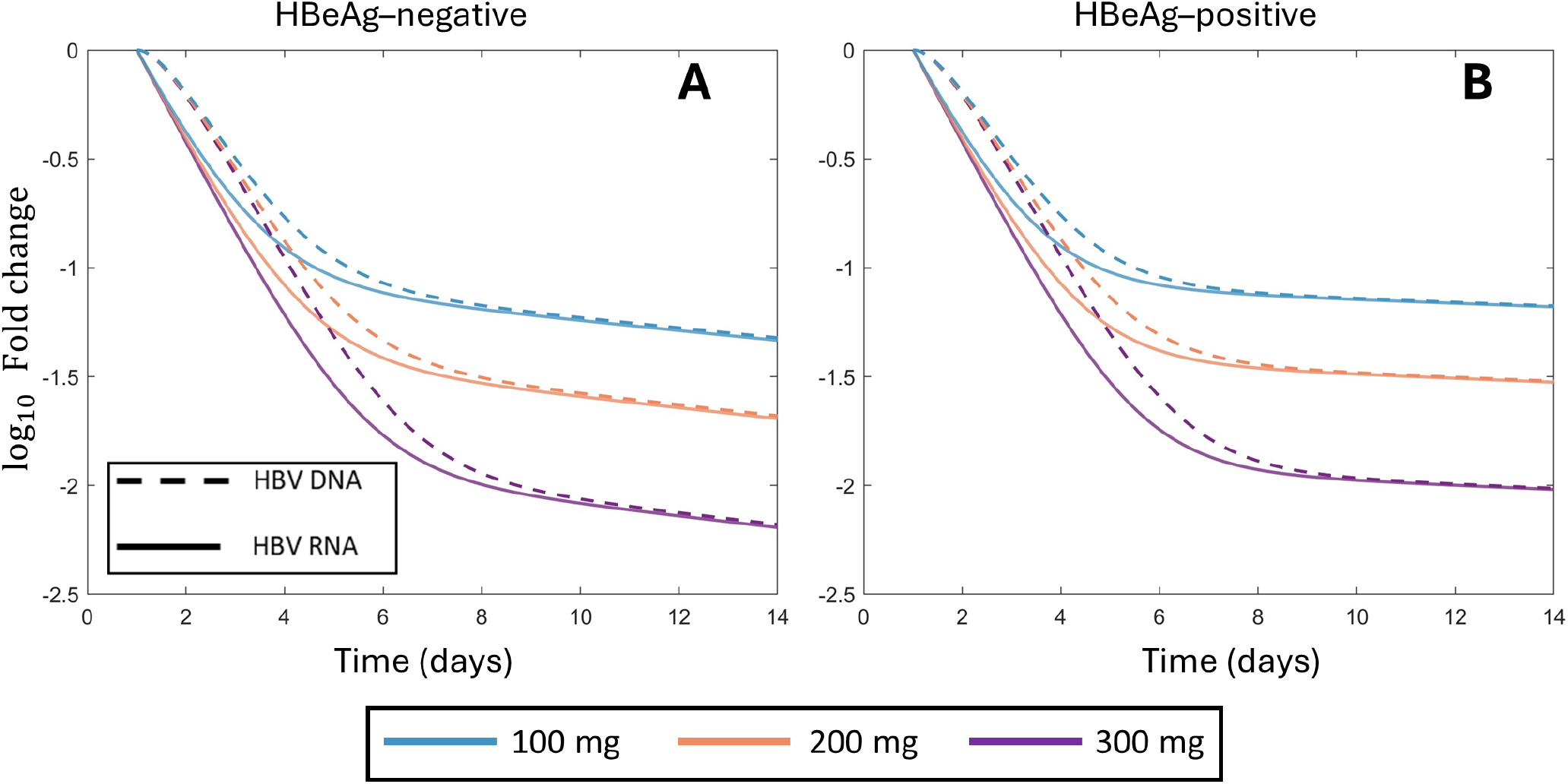
Larger fold decay in HBV RNA than HBV DNA during two weeks of treatment. Panels A and B show the fold decay in HBV RNA and HBV DNA during 14 days of treatment for 100 mg, 200 mg, and 300 mg of vebicorvir for the population parameter estimates for HBeAg-negative and positive participants, respectively.

In particular, the larger response of HBV RNA to changes in CAM effectiveness indicates that the dynamics of HBV RNA are more sensitive to treatment with a CAM than HBV DNA during the first phase of decline. As previously mentioned, this first phase of decline corresponds directly to the CAM mediated blocking of encapsidated pgRNA production. Following the initiation of treatment and on time-scales where infected cell death is negligible, intracellular encapsidated pgRNA and rcDNA amounts rapidly decay to a treated quasi-steady state. The crux of our analytical sensitivity analysis is tying the decay dynamics of these intracellular quantities to the extracellular dynamics of HBV RNA and HBV DNA. However, once these intracellular quantities reach their treated quasi-steady states, the dynamics of the HBV RNA and HBV DNA are similar. Consequently, it is not surprising that the differences in the dynamics of HBV RNA and HBV DNA are most pronounced during the first phase of treatment mediated decline. Indeed, at the first day post-treatment initiation for all three doses of vebicorvir, we calculate

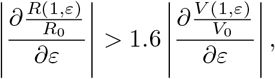

for both HBeAg-positive and negative participants. Accordingly, considering the early relative dynamics of HBV RNA may facilitate estimation of *ε* in on-going CAM monotherapy trials. However, the biological interpretation of our analytical result indicates that the utility of HBV RNA as a biomarker of CAM efficacy is limited to the first decay phase.

## Discussion

Many CAMs, including the first-generation agent, vebicorvir, have entered clinical trials and represent a promising treatment option for CHB. Here, we developed a multiscale model of CHB that bridges the intracellular viral life cycle and extracellular viral dynamics to understand the observed viral kinetics in a multiple ascending dose study of vebicorvir. Our model adapts the multiscale model of CHB developed by Goncalves et al. [36] to study the CAM RG7907 to include both the dynamics of ALT and uninfected hepatocytes. While multiscale models have been used in modeling chronic hepatitis C infection [22–24] and are beginning to be developed for CHB [36, 58–60], many previous modeling studies of CHB have not included the intracellular viral life cycle [29, 30, 43, 61–63]. Multiscale models, such as the model presented in this work or developed elsewhere [36, 59], offer unique insight into the intracellular and extracellular dynamics of HBV via the ability to explicitly model distinct mechanisms of action for novel small-molecule antiviral therapies.

We fit our model to the longitudinal HBV RNA, HBV DNA and ALT data of 29 individuals with chronic HBV infection treated with vebicorvir [4]. Our model fits the ALT, HBV RNA and HBV DNA dynamics well both during treatment and during follow-up. Vebicorvir treatment led to two observable phases of decline in HBV RNA and HBV DNA. The first phase of decline was rapid in both HBV RNA and HBV DNA, with a half-life of approximately 17 hours. Our analysis of the multiscale model suggests that this phase of decline is dominated by the clearance from the circulation of both HBV RNA and HBV DNA. Here, while our estimates for the clearance rate *c* are consistent with earlier results from Ribeiro et al. [29], they differ from the results of Goncalves et al. [36]. As mentioned, this difference is likely due to measurements of HBV RNA concentrations in the first week following treatment initiation, although Goncalves et al. [36] suggest that the pharmacokinetics of RG7907 may play a role. The second phase of decline of HBV RNA and HBV DNA was slower and our model analysis shows it is driven by the death of infected hepatocytes. We found that vebicorvir exhibits dose-dependent efficacy, with 300 mg daily dosing leading to the highest suppression of both HBV RNA and HBV DNA. However, unlike Goncalves et al. [36], who considered the CAM RG7907, our results do not indicate a HBeAg-dependent difference in drug efficacy. However, we identified significant HBeAg status dependent differences in the infection rate and death rate of infected hepatocytes, with higher values found in HBeAg-negative participants, possibly due to the loss of immune tolerance [55]. Despite the estimated higher infection rate in HBeAg-negative infection, we found that the significantly larger production rates of intracellular encapsidated pgRNA results in a larger basic reproduction number in HBeAg-positive infection. This finding is consistent with the higher viral load in HBeAg-positive infection.

Recently, there has been increased interest in using HBV RNA as a potential biomarker of treatment efficacy for CHB [35, 53]. As HBV RNA is a direct downstream product of cccDNA activity, via the production of encapisdated pgRNA, that is not directly impacted by treatment with NAs, decays in HBV RNA during NA treatment have been suggested to correspond to decays in cccDNA activity [35]. In particular, HBV RNA has been shown to predict viral rebound following treatment interruption in individuals treated with NAs [53]. Here, we evaluated HBV RNA as a biomarker of CAM efficacy. Specifically, we performed an analytic sensitivity analysis of our multiscale model and showed that HBV RNA concentrations are more sensitive to increases in CAM efficacy than HBV DNA concentrations. Our ability to distinguish between the HBV RNA and HBV DNA response to vebicorvir treatment crucially depends on our multiscale model explicitly including the dynamics of intracellular encapsidated pgRNA and rcDNA. Our analytical and simulation results suggests the continued use of HBV RNA as an important biomarker for CAM efficacy. Further, our modeling suggests that HBV RNA dynamics impart the most information regarding CAM efficacy during the first phase of decline. However, our modeling does not link cccDNA dynamics to observed changes in HBV RNA dynamics following CAM therapy.

Our modeling has some limitations. We did not include a mechanistic pharmacokinetic model to drive vebicorvir dynamics but rather assumed that vebicorvir concentrations rapidly reach steady-state concentrations during daily dosing. Consequently, we used a phenomenological model to capture vebicorvir washout and the resulting decline in CAM efficacy following treatment cessation. Using this model, we estimated a half-life of roughly 9 hours for vebicorvir, which is shorter than the estimated half-life of 23.5-28.4 hours observed by Yuen et al. [4]. Further, our multiscale model significantly simplified the intracellular life cycle and extracellular dynamics of HBV infection. At the intracellular level, we neglected cccDNA dynamics and potential rcDNA recycling within an infected cell in this short-term study, as the half-life of cccDNA has been estimated as approximately 40 days [64] and vebicorvir has not been shown to inhibit these processes [65]. However, as next-generation CAMs have demonstrated inhibition of rcDNA recycling, explicitly including cccDNA dynamics is a natural extension of our model. Furthermore, we did not model the dynamics of free pgRNA and other known HBV biomarkers, which are useful in diagnosing and treating CHB [66, 67], although an extension of our multiscale model could explicitly model the dynamics of these biomarkers.

All told, we developed a multiscale viral dynamics model to investigate the effect of vebicorvir monotherapy on the dynamics of HBV RNA and HBV DNA in chronically infected individuals. We note that, by including the intracellular dynamics of encapsidated pgRNA and rcDNA, our multiscale model can be used to study the effect of both NA and CAM plus NA combination therapy and future studies may include studying the effect of combination therapies on HBV dynamics. Here, we identified mechanistic differences between participants with HBeAg-positive and HBeAg-negative infection, showed that HBV RNA is more sensitive to CAM efficacy through an analytical study of our model, and finally predicted the time-scales on which HBV RNA dynamics are a potentially informative biomarker of CAM efficacy.

## Data availability

The viral load data used in this modeling study is available as a Excel file in the S2 Dataset. The Monolix model is available in Section *Monolix model used for data fitting* of S1 Text. The Matlab code used to generate Fig 6, and Fig A, B, C, D, E, F, and G of S1 Text is available at https://github.com/ttcassid/HBV_CAM_Treatment.

## Acknowledgement

We thank Assembly Biosciences for supplying the data used in this analysis.

## Supporting information

**S1 Text** Supporting information: Modeling resistance to the broadly neutralizing antibody PGT121 in people living with HIV-1

**S2 Dataset** Supporting information: Viral load data

